# Elaboration of the corticosteroid synthesis pathway in primates through a multi-step enzyme

**DOI:** 10.1101/2019.12.25.888461

**Authors:** Carrie F. Olson-Manning

## Abstract

Metabolic networks are complex cellular systems dependent on the interactions among, and regulation of, the enzymes in the network. However, the mechanisms that lead to the expansion of networks are not well understood. While gene duplication is a major driver of the expansion and functional evolution of metabolic networks, the effect and fate of retained duplicates in a network is poorly understood. Here, I study the evolution of an enzyme family that performs multiple subsequent enzymatic reactions in the corticosteroid pathway in primates to illuminate the mechanisms that shape network components following duplication. The products of the pathway (aldosterone, corticosterone, and cortisol) are steroid hormones that regulate metabolism and stress in tetrapods. These steroids are synthesized by a multi-step enzyme Cytochrome P450 11B (CYP11B) that performs subsequent steps on different carbon atoms of the steroid derivatives. Through ancestral state reconstruction and *in vitro* characterization, I find the ancestor of the CYP11B1 and CYP11B2 paralogs (in primates) had moderate ability to synthesize cortisol and aldosterone. Following duplication in the primate lineage the CYP11B1 homolog specialized on the production of cortisol while its paralog, CYP11B2, maintained its ability to perform multiple subsequent steps as in the ancestral pathway. Unlike CYP11B1, CYP11B2 could not specialize on the production of aldosterone because it is constrained to perform earlier steps in the corticosteroid synthesis pathway to achieve the final product aldosterone. These results suggest that pathway context, along with tissue-specific regulation, both play a role in shaping potential outcomes of metabolic network elaboration.

## Introduction

Biochemical networks consist of consecutive chemical reactions carried out by enzymes. In eukaryotes, the same pathway can be expressed across multiple tissues with each tissue possibly requiring different amounts of pathway products. Substitutions that lead to changes in enzyme function or expression that optimize pathway production in one tissue may have deleterious pleiotropic consequences in other tissues (Stern and Orgogozo 2008; Wray et al. 2003; Carroll 2005). To avoid these deleterious pleiotropic consequences, there are several mechanisms that allow for tissue specific regulation of widely expressed biochemical pathways: tissue-specific post-translational modifications (Ikegami et al. 2014), expression of different parts of the pathway in different tissues (Stearns 2010; Ramsay, Rieseberg, and Ritland 2009; Coberly and Rausher 2008), allosteric regulation where a metabolite (Snášel and Pichová 2019; Gui, Lewis, and Vander Heiden 2013) or another protein (Ikushiro, Kominami, and Takemori 1992; Nnamani et al. 2016; Perutz 1989) binds to an enzyme at a site other than its catalytic site affecting its activity.

Deleterious pleiotropic expression of a widely expressed pathway could be subverted with gene duplication followed by tissue-specific regulation and perhaps homology specialization. However, understanding how the mechanistic constraints of a pathway and its tissue-specific regulation changes following duplication requires an understanding of ancestral enzyme function, how these functions changed following duplication, and the effects of substitutions in those enzymes. Although there is much theory (Jensen 1976; Morowitz and Others 1999; Harborne 1990; Horowitz 1965) and comparative biology (Koes, Verweij, and Quattrocchio 2005; Chapman and Ragan 1980; Jensen 1985) on how biochemical pathways are built and change, to date, there is little empirical evidence on biochemical pathways change on evolutionary timescales.

The corticosteroid synthesis pathway in vertebrates provides a model to understand how a biochemical pathway resolves a duplication of one of its components that result in tissue-specific homologs with divergent functions. The corticosteroids are tetrapod hormones that regulate metabolism and stress. The corticosteroid biosynthetic pathway (Fig 1A) is a well-studied example of a pathway that uses a variety of regulatory mechanisms to produce tissue-specific expression of its products across different tissues (Vinson 2016). The enzyme in the adrenal cortex that synthesizes corticosteroids is Cytochrome P450 11B (CYP11B) (Fig 1B). CYP11B is a multi-step enzyme in that it is a single enzyme that performs distinct and consecutive reactions of a biochemical pathway. CYP11B first modifies the 11-carbon atom of the steroid precursors 11-deoxycorticosterone (DOC) and 11-deoxycortisol and subsequently oxidizes the 18-carbon of corticosterone to produce aldosterone (Kominami, Harada, and Takemori 1994).

**Figure 1.**
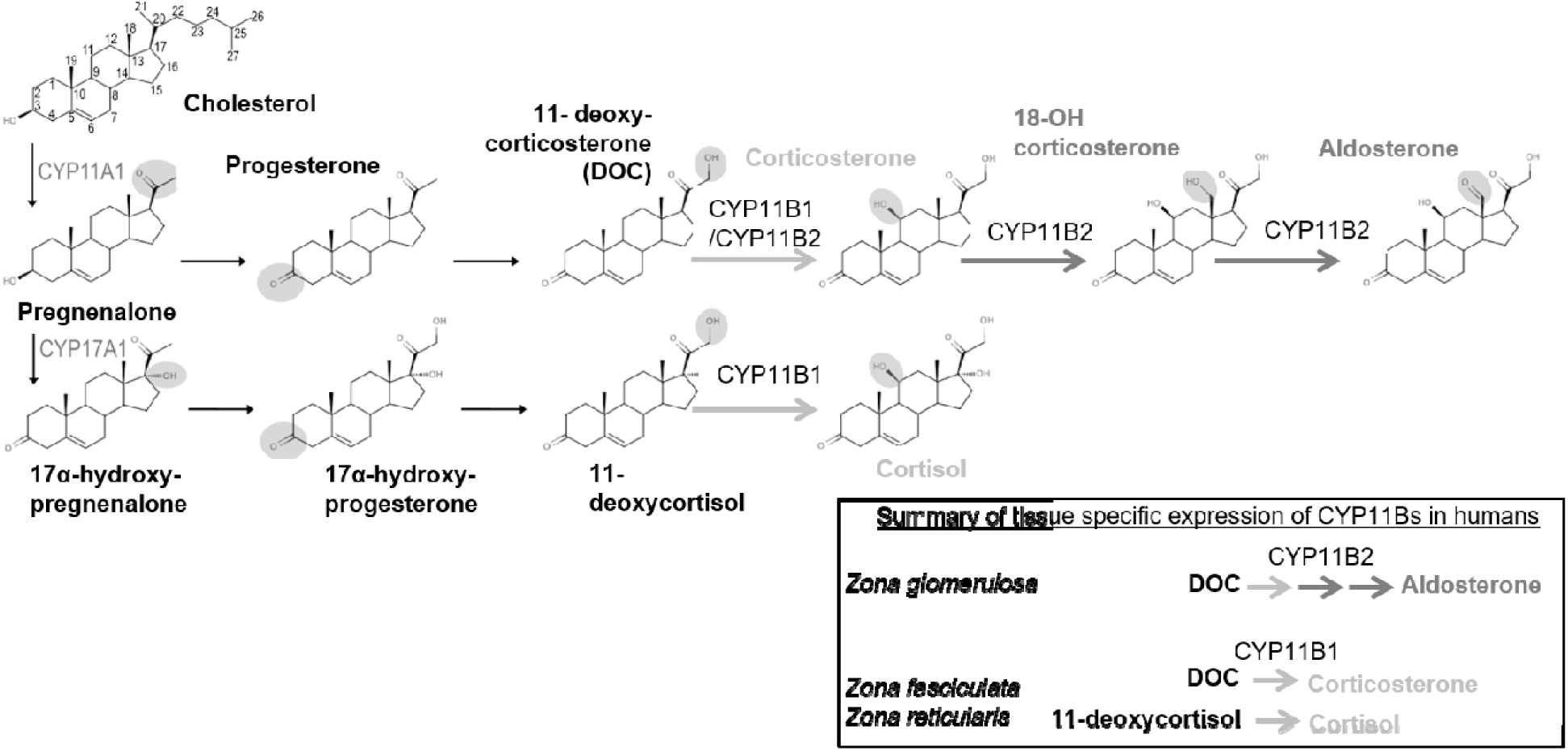
Synthesis of corticosteroids in humans **(A)** is accomplished by the CYP11B homologs. CYP11B1 and CYP11B2 hydroxylate the 11-carbon of 11-deoxycorticosterone (DOC) or 11-deoxycortisol (light grey arrows) to produce corticosterone and cortisol. After performing 11β-hydroxylation on 11-deoxycorticosterone, CYP11B2 oxidizes the 18-carbon of corticosterone and 18-hydroxycorticosterone (dark grey arrows) to produce aldosterone. In humans, the two paralogs of CYP11B are expressed in different layers of the adrenal cortex **(inset)**. Cortisol and corticosterone (produced by 11-carbon hydroxylation by CYP11B1) are preferentially produced in the *zona fasciculata* and *zona reticularis* of the adrenal cortex. Aldosterone is preferentially produced by CYP11B2 in the *zona glomerulosa* by performing both 11-carbon hydroxylation of 11-deoxycorticosterone and 18-carbon oxidation. Other Cytochrome P450s in the corticosteroid pathway are labeled and new modifications for each enzymatic step are shaded.

In all tetrapods studies thus far, 18-oxidized steroids and 11-hydroxylated steroids are produced in different tissues of the adrenal cortex. Aldosterone is produced in the zona glomerulosa and the 11β-corticosteroids (cortisol and corticosterone) are produced in the zona reticularis and zona fasciculata (Fig 1B). In addition to producing different steroid hormones, the zona have different cellular and topological morphologies (Vinson 2016). This zona specific synthesis (zonation) of corticosteroids persists in species with one CYP11B enzyme (Sanders et al. 2016; Boon et al. 1997; Schiffer, Anderko, and Hannemann 2014; Nonaka et al. 1995) or those that have experienced duplications and have two CYP11B homologs (Nishimoto et al. 2010; Domalik et al. 1991; Ogishima et al., n.d.; Malee and Mellon 1991; Miller and Auchus 2011; Payne and Hales, n.d.; Lisurek and Bernhardt 2004; Bassett, White, and Rainey 2004). Previous work on the CYP11B homologs in primates suggests that following a duplication, the CYP11B paralogs have specialized on either 18-oxidation or 11-hydroxylation (Zöllner et al. 2008; Hobler et al. 2012; Strushkevich et al. 2013), but none have taken a historical approach (Harms and Thornton 2013) to understand how function changed in this pathway and the mechanisms that have shaped that change.

The mechanisms that allow the production of different metabolites in the different tissues of the adrenal cortex in single-CYP11B species are unclear but likely causes include tissue-specific expression of other pathway members (Nishimoto et al. 2010), allosteric regulation of CYP11B by other enzymes (Ikushiro, Kominami, and Takemori 1992), or the tissue-specific expression of CYP17A (Figure 1). In species with two CYP11Bs (some primates and, independently, some rodents (Baker, Nelson, and Studer 2015)), zonation is controlled by a combination of tissue-specific expression of the paralogs (Nishimoto et al. 2010; Domalik et al. 1991; Ogishima et al., n.d.; Malee and Mellon 1991) and enzyme substrate specialization. In humans, for example, CYP11B1 is expressed in the zona reticularis and zona fasciculata and synthesizes the 11β-oxidized cortisol and corticosterone (Zöllner et al. 2008). CYP11B2 produces the 18-oxidized corticosteroid aldosterone (Hobler et al. 2012) and is expressed only in the zona glomerulosa.

In all accounts, it appears that the physiological function and, in particular, the zonation of the corticosteroids in the adrenal cortex has been maintained in mammals with one or two CYP11B enzymes. The misregulation of these enzymes is catastrophic. Altered cortisol levels in humans leads to metabolic syndrome, mood disorders, and bone loss (Marques, Silverman, and Sternberg 2009) and the misregulation of aldosterone leads to hyper- or hypo-tension, hypo-kalmia and androgen excess (Connell, Fraser, and Davies 2001; Peter, Dubuis, and Sippell 1999). Therefore, following the duplication of CYP11B, it is critical that the levels of corticosteroids produced are properly balanced as to avoid any deleterious pleiotropic effects.

In this study, I dissect the changes to the intrinsic biochemical functions of CYP11B paralogs following the duplication in the corticosteroid pathway in simian primates (infraorder Simiiformes) from their common ancestor. I focus our study on the *in vitro* functional changes in the primate lineage. To do so I characterized the specific activity prior to and after its duplication in the primate lineage, with emphasis on CYP11B2 enzyme (sometimes called aldosterone synthase), including a detailed dissection of the trajectory of evolutionary change in the CYP11B2 lineage. Following duplication of the CYP11B multifunctional enzyme, both 11- and 18-oxidation were maintained in tissues where 18-oxidation is required. These findings have implications for the elaboration of biochemical pathways and on constraints on specialization when the enzymes are multi-functional.

## Methods

### Phylogenetics and ancestral state reconstruction

We sampled 86 sequences in tetrapods, cartilaginous and bony fishes from NCBI, Uniprot, and Ensembl. I aligned the sequences first with MUSCLE (Edgar 2004), and manually checked for alignment errors and removed species-specific insertions. With PhyML (Guindon and Gascuel 2003, Guindon et al. 2010) and SPR+NNI rearrangements to infer the maximum likelihood (ML) phylogeny of our vertebrate CYP11B/C sequences under the JTT+G+I model, which was the best-fit model as evaluated by PROTTEST (Darriba et al. 2011) using AIC. Insertions and deletions were resolved with maximum parsimony (Felsenstein 2003).

There were minor incongruences between the ML phylogeny and the known mammalian species phylogeny (Sen Song et al. 2012) that led to two hypotheses about the evolution of CYP11B. A more parsimonious explanation of gene gains and losses places the duplication of CYP11B in the Old World primates (Figure S2), and the placement of the New World primate orthologs within the Old World primate CYP11B1 clade in the ML topology is incorrect. I reconstructed the ancestral states with (Yang 2007) implemented in Lazarus (Hanson-Smith, Kolaczkowski, and Thornton 2010) for both topologies. The reconstructed sequences for the ancestor of all Old World primate CYP11Bs (which is also ancestral to New World primate CYP11Bs in the ML topology), the ancestor of primate CYP11Bs, and the ancestor of primate CYP11B2s are identical between topologies.

To test whether our functional inferences are robust to statistical uncertainty in the reconstruction, I made an alternative ancestor for the ancestral Old World primate CYP11B that included one site (A438T) in the reconstruction that had an alternate state with a posterior probability greater than 0.2.

### Plasmid construction

Unlike (Hobler et al. 2012; Zöllner et al. 2008) I did not get high expression of the human sequence of CYP11B1 or CYP11B2 in *Escherichia coli* under published expression conditions. Instead, I ordered sequences that were codon optimized for expression in *E. coli* as IDT gblocks (Integrated DNA Technologies). The *E. coli* codon optimized human cDNA and the inferred ancestral CYP11B cDNAs were expressed in the plasmid Novagene pET17b plasmid cloned with the restriction sites NdeI/HindIII as in (Hobler et al. 2012) with slight modifications. Briefly, the mature form of the protein was expressed without the N-termini mitochondrial targeting sequence. The new N-terminus was modified from GTRAAR to MATKAAR (Hobler et al. 2012) and a hexa-histidine tag added on the C-terminus to assist with purification (Nonaka et al. 1992). Individual point mutations were introduced via site-directed mutagenesis.

The mature human ferrodoxin 1 (FdX) was expressed in pET17b was amplified as in (Uhlmann et al. 1992) as a cytosolic protein. The mature peptide (sites 61-184 beginning as MSSSEDKI) was amplified with the restriction sites NheI and HindIII and cloned into the pET17b vector. The mature human FdxR (beginning as MSTQEK (Sagara et al. 1993)) was amplified with restriction sites NdeI and KpnI and cloned into the vector pET17b. The chaperones GroEL and GroES (Nishihara et al. 1998) were obtained in the pGRO12 plasmid (from Dr. Michael Waterman).

### Protein expression CYP11B

pET17b-CYP11B (AmpR) plasmids were co-transformed with the pGRO12 plasmid (KanR) into competent BL21(DE3) *E. coli* cells (Zöllner et al. 2008). All P450-expression cell lines were grown in terrific broth (TB) supplemented with 50 ug/ml ampicillin and 50 ug/ml kanamycin. For expression, a scraping of frozen glycerol stock was inoculated into 5 ml TB with the same amount of antibiotics under shaking (225 rpm) overnight at 37 C. I had very poor expression when overnight cultures were started from anything other than a glycerol stock or if the antibiotic was carbenicillin instead of ampicillin. 4 ml of the overnight culture was added to 400 ml TB (ampicillin, kanamycin) in a 2.8 L baffled flask and shaken at 210 rpm at 37 C. When the optical density at 600 nm (OD_600_) reached 0.7, I added 1 mM **δ**-aminolevulinic acid, 0.5 mM IPTG, and 4 mg/ml arabinose and changed the incubation parameters to 27.5 C and 76 rpm. Four hours later, 10 ul of 25 mg/ml chloramphenicol was added and shaking was changed to 125 rpm. Shaking continued for 16 hours before harvest.

Cells were harvested under 4 C, 4000 *x g* for 30 minutes. Pellets were frozen at −80 C and resuspend in CYP11B buffer A (50 mM potassium phosphate (pH 7.4), 0.1 mM EDTA, 20% glycerol, 1.5% sodium cholate, 1.5% Tween 20, 0.1 mM PMSF, 0.1 mM DTT) with sonication. The sonication conditions were: setting 8 on the Q55 Sonicator (QSonica), 15 second on, 15 seconds off four times. After resting on ice for one minute, the sonication protocol was repeated three times.

After sonication the cell debris was removed by centrifugation at 19,000 *x g* for 45 minutes. The supernatant was passed through a 0.45 micron low-protein binding filter, applied to a 5 ml Ni-NTA agarose column bed, and equilibrated with CYP11B Buffer A with 40 mM imidazole. Proteins were eluted with CYP11B buffer B (200 mm imidazole acetate, pH 7.4, 20% glycerol, 0.1 mm EDTA, 0.1 mm dithiothreitol, 1% sodium cholate, 1% Tween 20). Following elution, fractions containing the P450 (red fractions) were combined and concentrated in concentrating tubes with a 30,000 kDa membrane at 4 C. After one wash with CYP11B buffer A, the proteins were concentrated in a Slide-A-Lyzer dialysis cassette (Thermo Scientific) with the concentrating solution of 40% PEG 20 (Sigma-Aldrich Product No. 81300) at 4 C for up to 2 hours. Aliquots of concentrated enzyme were flash frozen in liquid nitrogen and stored at −80 C. Frozen samples maintained activity for at least six months.

The concentration of active enzyme was calculated with the ferrous carbon monoxide (CO) versus ferrous difference spectrum (Guengerich et al. 2009). I added 10-20 μl of concentrated P450 to CO buffer (100 mM potassium phosphate buffer (pH 7.4), 1 mM EDTA, 20% glycerol, 0.5% sodium cholate, and 0.4% Triton N-1010) to a total volume of 1 ml. After blanking with the P450 and buffer solution, CO was bubbled into the cuvette at 2 bubbles per second for 45 seconds. A few milligrams of sodium dithionite powder is added to reduce the CO-P450 complex and the cuvette was covered with parafilm and inverted ten times. In a cuvette in the Nanodrop 2000c I scanned the spectra from 400 to 500 nm every thirty seconds for up to 15 minutes, when the spectra reached equilibrium, or when the difference between absorbance at 450 nm and 490 nm was the greatest (Omura and Sato 1964). I calculated the total amount of active enzyme by taking the difference of the absorbance at 450 nm from 490 nm with an extinction coefficient of P450 0.091 mM-1*cm-1 [(Δ*A*_*450*_-Δ*A*_*490*_)/0.091 = ηmol of P450 per ml].

### Protein expression Reductases

The BL21(DE3) cells with FdX were grown overnight (100 ug/ml AMP in LB) and 5 ml of overnight culture was added to 500 ml LB with no antibiotic at 37 C 210 rpm. When the OD_600_ was 0.7, the cells were induced with 0.5 mM IPTG and grown at 30 C, 125 rpm for 24 hours (Modified from (Uhlmann et al. 1992)).

FdX containing cells were harvested via centrifugation 4000 *x g* for 25 minutes at 4 C. The pellet was resuspended in 30 ml Buffer P (10 mM KPO4 pH 7.4, 1 mM EDTA, 1 mM PMSF, 30 ul lysozyme and DNaseI) from (Uhlmann et al. 1992) and homogenized with sonication as outlined above for the P450 enzymes. The large cell debris was removed by centrifugation at 10,000 *x g* for 30 minutes at 4 C. AdX was enriched by adding ammonium sulfate to 60% at 4 C followed by centrifugation at 45,995 g for 1 hour. The supernatant was dialyzed overnight with Buffer P. The supernatant was passed through a 0.45μm filter, added to Source Q anion exchange column (GE Healthcare Life Sciences) and eluted with a 0-400 mM NaCl gradient in Buffer P. The brown fractions were concentrated and stored in 20% glycerol Buffer P. and quantified in a nanodrop cuvette with the extinction coefficient of 9.8 mM^−1^cM^−1^ at 415 nm.

The human FdxR was also expressed in pET17b and was co-expressed with GroEL/ES in BL21(DE3). Cells were grown overnight (37 C, 225 rpm) and inoculated into 500 ml TB with 100 μg/ml ampicillin and 50 μg/ml kanamycin. At OD_600_ of 0.5 FdxR was induced with 0.5 mM IPTG and 4 mg/ml arabinose. After induction, the cells were grown at 26 C, 125 rpm shaking for 72 hours (modified from (Sagara et al. 1993)).

Cells were harvested via centrifugation 4000 *x g* for 25 minutes at 4C and resuspend in 30 ml Buffer P (10 mM KPO4 pH 7.4, 1 mM EDTA, 1 mM PMSF, 30 ul lysozyme and DNaseI). The cells were homogenized via sonication as described above. The cell debris was removed by centrifugation at 45,995 *x g* for 1 hour at 4 C. The supernatant was dialyzed overnight with Buffer P, and applied to a Source Q anion exchange column after filtration through a 0.45 micron filter. The flow through was collected and applied to a 2’5’-ADP-Sepharose column. The column was washed with Buffer P and the protein eluted with a 0-400 mM NaCl gradient in Buffer P. The flow through on the 2’5’-ADP Sepharose column was reapplied to obtain more FdxR. Yellow fractions were collected, concentrated and stored in 20% glycerol Buffer P. FdxR was quantified using the extinction coefficient 10.9 mM^−1^cM^−1^ at 450 nm in Buffer P.

### Cytochrome P450 kinetic assays

To quantify the specific activity (μmol min^−1^ mg^−1^) of the CYP11B homologs, I combined 0.4 μM enzyme with 50 uM HEPES, 1 μM MgCl_2_, 0.5 mM FdxR, 10 mM FdX, 5 μM glucose-6-phosphate, 0.1 units glucose-6-phosphate dehydrogenase (Sigma), and 350 μM substrate in 100 μL reaction volume. The reactions were placed at 37 C for 3 minutes and the reaction was started by adding NADPH to 2 mM. The reaction was incubated at 37 C with slight agitation for 15 minutes and stopped with 150 μL chloroform. After the reaction was stopped, I added cortisol to 100 μM as an internal standard. The steroids were extracted three times with 100 μL chloroform and dried in a hood. After the chloroform dried, I added 20% HPLC-grade acetonitrile to resuspend the steroids (modified from (Hobler et al. 2012)).

All steroids (products and substrates) were monitored with LC-MS methods. The samples were run on an Agilent 1200 Series HPLC coupled to an Agilent 6410 triple quadrupole mass spectrometer (Morton, Roux, and Bergelson 2015). 1 μl of sample was injected onto a 4.6 × 50-mm Agilent ZORBAX Eclipse Rapid Resolution HT XBD-C18 column with a 1.8 μM particle size. All samples were run with the following program: 0.3 mL/min flow rate, ramp from 60:40 0.1% formic acid in HPLC-grade water:acetonitrile to 30:70 ratio over ten minutes. The mass spectrometer was run in precursor positive-ion electrospray mode, monitoring all parent ions (Table S2). The fragmentor voltage was optimized with steroid standards for aldosterone, cortisol, corticosterone, and 11-deoxycorticosterone (DOC) obtained from Sigma-Aldrich and Steraloids. Maximum detection was achieved with collision energy (Table S2). Individual steroids were identified based on comparison to purchased standards, fragmentation pattern, and retention time as compared with a cortisol internal standard. The final data were generated by dividing the parent ion counts by the internal cortisol standard.

## Results

### CYP11B duplicated in the primates

We studied a duplication followed by functional divergence in the corticosteroid pathway in primates. As previously reported (Baker, Nelson, and Studer 2015), I found at least three independent duplications of CYP11B in mammals (Figure S1). Here I focus on the duplication and subsequent functional shift that occurred in the primate lineage (Figure 2). The phylogenetic evidence is nearly equivocal on whether the gene duplication occurred before or after the split of Old World primates (parvorder Catarrhini) and New World monkeys (parvorder Platyrrhini), with the ML tree weakly supporting the duplication occurring before this split, and a more parsimonious hypothesis of gene gain and loss favoring the duplication occurring after this split.

**Figure 2.**
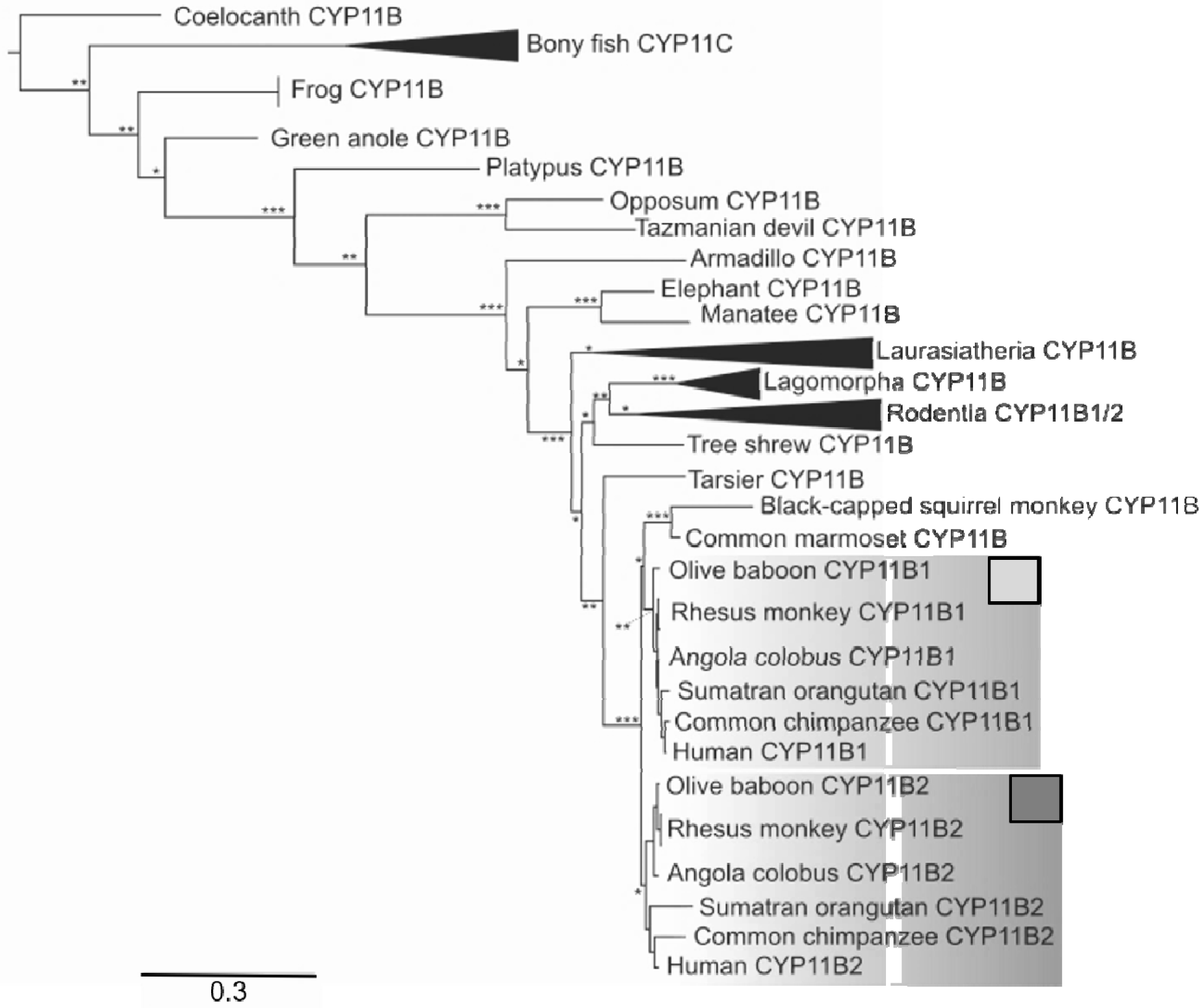
Maximum likelihood gene phylogeny of 86 vertebrate CYP11B/C sequences with evolutionary model JTT+I+G (full maximum likelihood gene phylogeny in Figure S1). The clades of bony fish CYP11C, Laurasiatheria (including bats, carnivorans, whales and even-toed ungulates), Lagomorpha (rabbit and pika) and Rodentia are collapsed for clarity. The Old World primate phylogeny is recapitulated twice on the phylogeny with the CYP11B1 clade (light grey box) and the CYP11B2 clade (dark grey small box). The approximate likelihood statistic indicates node support at each node is indicated with asterisk (*>4, **>10, ***>50). The reconstructed ancestors of the CYP11B1, CYP11B2, and the node of those clades which is the common ancestor of all primates in the infraorder Simiiformes.

Of the New World monkeys for which I found sequences, the black-capped squirrel monkey (*Saimiri boliviensis boliviensis)* had two copies of CYP11B. However, this apparent second duplication is species-specific and occurred after the divergence of New and Old World primates. The closest relatives of New and Old World primates (Sen Song et al. 2012), including the Chinese tree shrew (*Tupaia chinensis*) and tarsier (*Tarsius syrichta*) have a single copy of CYP11B. I reconstructed the duplication node and the nodes that led to the two CYP11B paralogs for both phylogenies.

### Human CYP11B1 and CYP11B2 functional divergence

The CYP11B paralogs in humans have functionally diverged following duplication. CYP11B increased its ability to produce either cortisol and corticosterone (11β-hydroxylation products) and most of its ability to produce aldosterone (Figure 3) (consistent with previous studies (Zöllner et al. 2008; Hobler et al. 2012; Strushkevich et al. 2013)). These results agree with these previous studies except that I detected a small amount of 18-hydroxycorticosterone and aldosterone produced by CYP11B1 *in vitro* (Figure 3).

**Figure 3.**
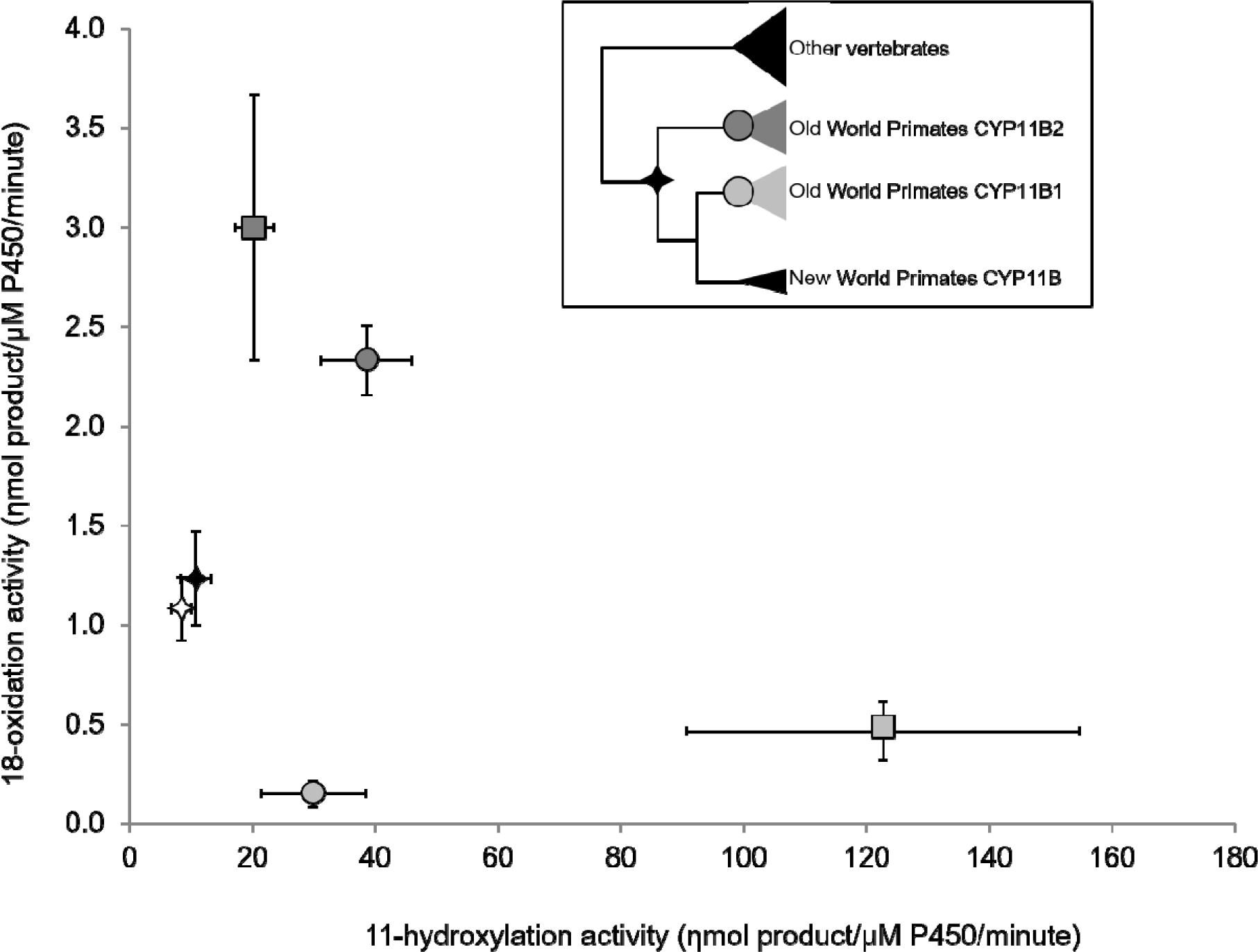
The primate AncCYP11B (black star) and the alternate ancestral sequence (white star) had moderate 11β-hydroxylation and 18-oxidation while the other paralogs have specialized. The inset depicts a reduced maximum likelihood phylogeny of CYP11B evolution in primates. The error bars indicate the 95% confidence intervals. The extant human paralogs CYP11B1 (light grey square) and CYP11B2 (dark grey square) have increased 11-hydroxylation and 18-oxidation activities. The intermediate ancestor, AncCYP11B1 (light grey circle), has lost most of its 18-oxidation activity and AncCYP11B2 (dark grey circle) has increased both activities.

The human CYP11B2 paralog has increased ability to produce both 11β-hydroxylation products and 18-oxidation products (aldosterone) (Figure 3). 11β-hydroxylation function is always higher than 18-oxidation function and both paralogs retain the ability to complete both 11β- and 18-oxidation.

### Function of ancestral CYP11B, CYP11B1, and CYP11B2

The pre-duplication ancestor of CYP11B1 and CYP11B2, AncCYP11B, had moderate 18-oxidation activity and relatively low 11β-hydroxylation activity *in vitro* (Fig 3). This result is robust to statistical ambiguity in the reconstruction: an alternate reconstruction incorporating all second-best states with PP > 0.2 (AncCYP11B-AltALL) has indistinguishable function from the most probable ancestor. Both of the AncCYP11B1 and AncCYP11B2 reconstruction have an average posterior probability greater than 0.99 and have no sites with an alternate reconstruction with a posterior probability higher than 0.2.

AncCYP11B1 and AncCYP11B2 enzymes have differentiated from the AncCYP11B toward 11β- or 11-oxidation and 18-oxidation activity, respectively (Fig 3). With respect to AncCYP11B, AncCYP11B1 had a moderate increase in 11β-hydroxylation function and a decrease in 18-oxidation function. Between AncCYP11B1 and human CYP11B1, there was a 4-fold increase in 11β-hydroxylation activity. AncCYP11B2 had a two-and-a-half and four-fold increase in 18-oxidation and 11β-hydroxylation, respectively. From AncCYP11B2 to human CYP11B2, there was only a slight increase in 18-oxidation and a slight decrease in 11β-hydroxylation. The AncCYP11B2 and the extant human CYP11B2 both maintain 11-hydroxylation and 18-oxidation activity.

However, it is important to note that CYP11B2 did not specialize in the traditional sense in that it did not lose one function in favor of another. It appears that it is necessary that in order to increase 18-oxidation activity, there must also be an increase in 11β-hydroxylation activity. Therefore, I dissected the substitutions from the preduplicated AncCYP11B to AncCYP11B2 to understand the interplay of balancing 18-oxidation and 11β-hydroxylation activities.

### Substitutions leading to AncCYP11B2

To understand the trajectory of functional evolution in the CYP11B2 lineage, I characterized the function of all intermediates from AncCYP11B to AncCYP11B2. There are four substitutions along this lineage. I dissected the effect of those substitutions by characterizing the 11β-hydroxylase (Fig 4A) and 18-oxidation (Fig 4B) functions of all 16 combinations. All four substitutions appear to affect either 11β-hydroxylation or 18-oxidation activity (Fig 4). Here I depict ancestral character states as lowercase and derived as uppercase letters with the amino acid position between; the four substitutions are d83N, d302E, q472L, and s492G.

**Figure 4.**
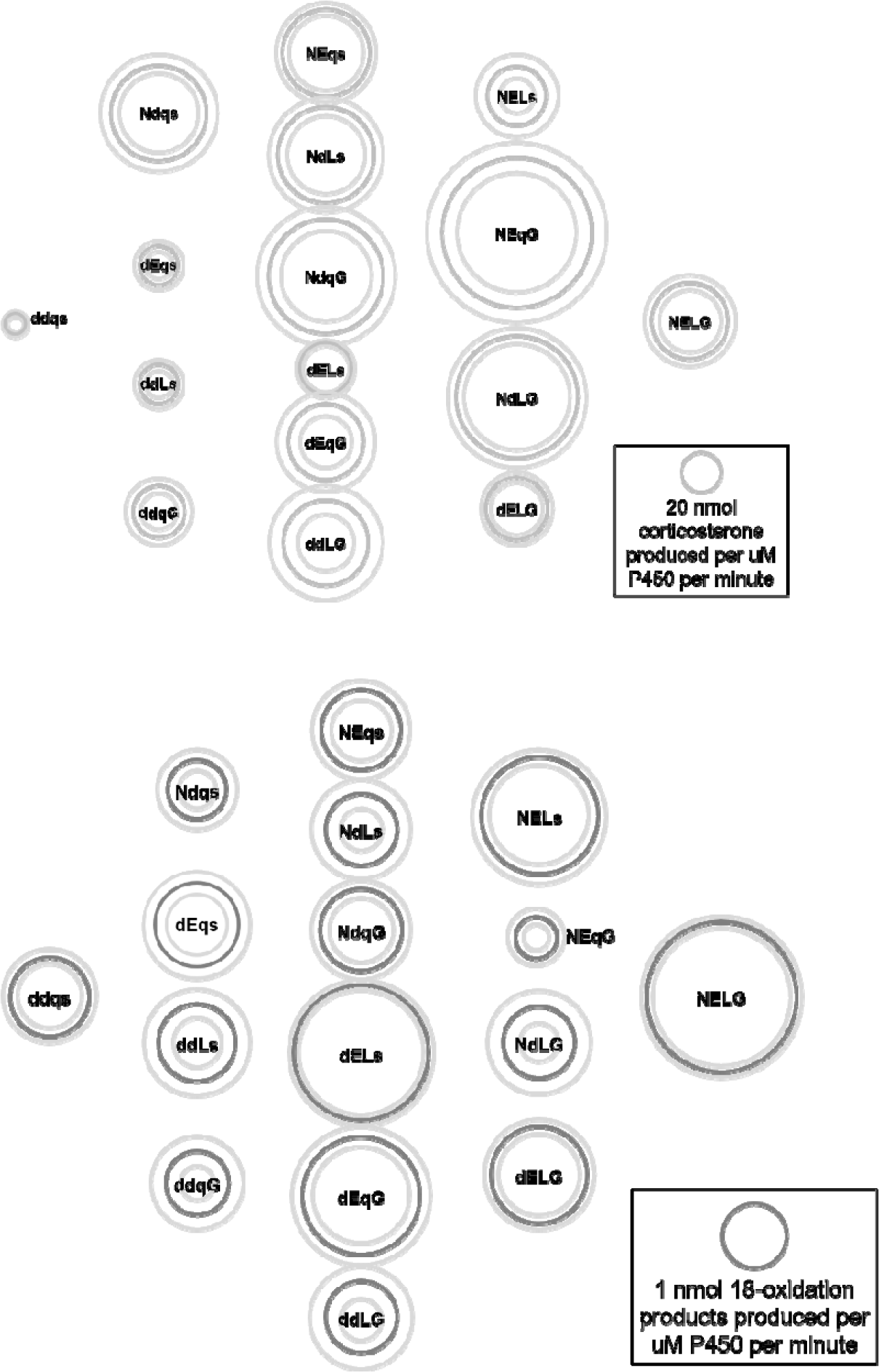
Genetic dissection of 11β-hydroxylation (A) and 18-oxidation (B) activity in the CYP11B2 lineage from AncCYP11B to AncCYP11B2. The genotypes are depicted in or near the circles with lowercase indicating ancestral and uppercase as derived. Moving from left to right, single derived substitutions are added. The diameter of dark circles indicates the mean activity in nmol product/uM P450/minute for that genotype as indicated by the key in the box. The light grey circles surrounding the mean specify 95% confidence intervals.

When I dissect the intrinsic function of all possible intermediates, Two of the substitutions (d83N and s492G) always increase 11β-hydroxylation function in any genetic background tested (Table 1, Fig 5A). Consistent with this observation, on the trajectory between AncCYP11B and AncCYP11B2, only genotypes with both 82N and 492G (NEqG, NdLG, NdqG, and NELG) have the highest 11β-hydroxylation activity. The other two substitutions (d302E and q472L) have mixed effects possibly caused by non-significant background-dependent effects (Table 2, Fig 5B). None of the substitution combinations lead to complete loss of either function, and only one genotype (NEqG) has 18-oxidation activity that is lower than the ancestral AncCYP11B genotype (ddqs). Therefore, most mutational trajectories are plausible between AncCYP11B and AncCYP11B2, except maybe through NEqG.

**Table 1:** Average effect of adding derived substitution.

| Substitution | 11β-hydroxylation<br>(nmol/uM450/min) | 18-oxidation<br>(nmol/uM450/min) |
| --- | --- | --- |
| d82N | 23.62 ± 9.13 | -0.35 ± 0.29 |
| d302E | -3.65 ± 9.97 | 0.38 ± 0.39 |
| q472L | -0.47 ± 11.73 | 0.10 ± 0.28 |
| s492G | 18.07 ± 6.45 | -0.19 ± 0.36 |

**Table 2:** Average effect of completing epistatic interaction.

| Comparison | 18-oxidation<br>(nmol/uM450/min) |
| --- | --- |
| xdLx to xELx | 0.48 ± 0.57 |
| xdxx (not dL) to xExx (not EL) | 0.16 ± 0.56 |
| xEqx to xELx | 0.29 ± 0.66 |
| xxqx (not Eq) to xxLx (not EL) | -0.12 ± 0.18 |

**Figure 5.**
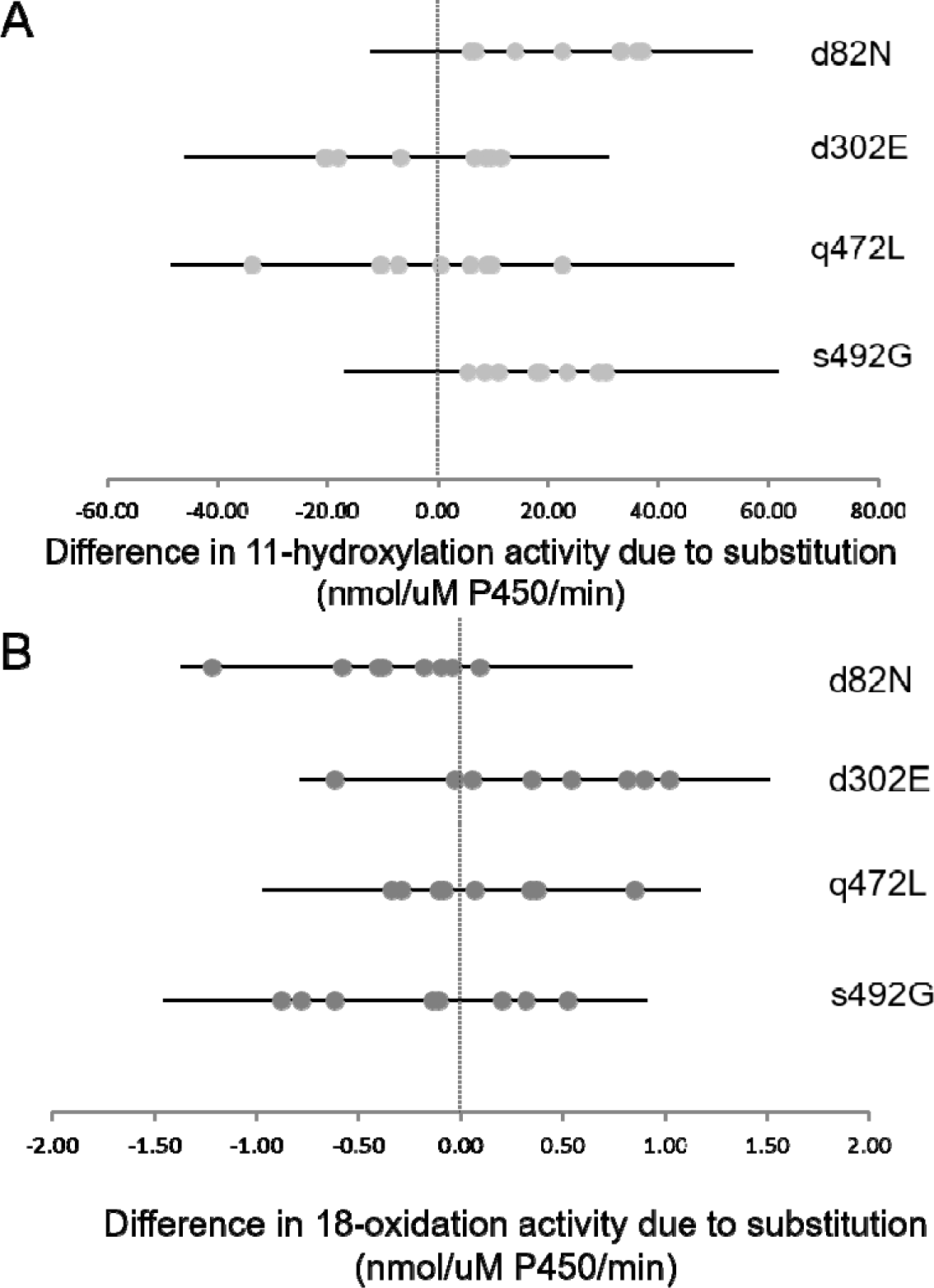
The distribution of functional effect of each substitution across all tested backgrounds, calculated by subtracting derived from ancestral activity. 11β-hydroxylation activity (A, light grey) increases in every background for the d83N and s492G substitutions. 18-oxidation activity is partially controlled by non-significant background-dependent effects d302E and q472L (B, dark grey) and its pattern is more complicated.

Structural analysis does not provide much insight but is reported nonetheless. I mapped the substitutions on the human CYP11B2 crystal structure (PDB: 4dvq (Strushkevich et al. 2013)) and found that none of the substitutions are in the canonical substrate recognition regions (Gotoh 1992) or even within 12 Angstroms of the heme or substrate. Any structural modifications must have subtle effects on the substrate recognition or binding regions.

## Discussion

Here I studied the evolution of the corticosteroid synthesis pathway to determine how functional evolution is resolved for a multi-step enzyme after duplication. The corticosteroids are a crucial pathway, the products of which control metabolism, salt balance, and immunity among other processes. To understand the processes that shape the augmentation of this pathway (two enzymes that do the job of one), I used ancestral state reconstruction to resurrect enzymes bracketing a gene duplication in the corticosteroid synthesis pathway in primates. Prior to its duplication, the AncCYP11B enzyme had moderate activity on both the 11β- or 18-carbon of the precursor DOC. Following duplication, while the CYP11B1 paralog specialized on 11β-carbon activity, CYP11B2 maintained multiple functions, similar to the that of AncCYP11B, but increased.

As biochemical pathways evolve, both mechanistic enzymatic and tissue-specific constraints shape their evolution. Previous biochemical studies have established that CYP11B 11β-hydroxylation activity occurs first followed by 18-oxidation (Kominami, Harada, and Takemori 1994). However, if the 11β-hydroxylated substrate (i.e. corticosterone) leaves the active site, it cannot reenter to be 18-oxidized to aldosterone. This implies that in order to have tissue-specific expression of CYP11B duplicates and still maintain zonation, CYP11B2 must maintain both functions rather than specializing on 18-oxdiation activity. Our results that consider the ancestral CYP11B activity help explain why CYP11B2 is constrained to maintain 11-hydroxylation activity in addition to 18-oxidation even when the conserved function of the zG is to produce 18-hydroxylated products.

The resolution of multi-functional (more than one defined function on preferred substrates) or promiscuous (trace activity for a non-preferred substrate) enzymes have been well studied (Huang et al. 2012; Voordeckers et al. 2012; Thomson et al. 2005; Des Marais and Rausher 2008; Deng et al. 2010; Boucher et al. 2014) and the prevailing model of functional evolution following a duplication in most of these studies is the escape from adaptive conflict model (with (Boucher et al. 2014) finding a rare case of neofunctionalization). However, under the umbrella of multi-functional enzymes are multi-step enzymes. Multi-step enzymes, like CYP11B, provide an understudied avenue in which biochemical pathways change their configuration without gaining a novel synthetic step. CYP11B1 underwent classic duplication, degeneration, and complementation (DDC) model where one function was maintained (Force et al. 1999). However, none of the classic models fit for CYP11B2 as both functions were maintained, if not slightly improved following duplication. When the complexity of biochemical mechanisms of metabolism are considered with the further complication of tissue-specific regulation, further models may be required, including one to include the maintenance of function in enzymes that are themselves short biochemical pathways like CYP11B. It is yet unknown how common or important such multi-step enzymes are in metabolism.

We tested the intrinsic activities of these enzymes *in vitro*, but predict that some of their *in vivo* activities may differ depending on the other enzymes with which they are expressed in the tissues of the adrenal cortex. For example, the activity of CYP11B in bovine (which has a single CYP11B copy - similar to state in the ancestral primate) is allosterically regulated by CYP11A *in vitro* (Ikushiro, Kominami, and Takemori 1992). This may partially explain how the adrenal cortex in bovine still expresses zonation of cortisol and aldosterone production with only a single copy of CYP11B. When co-expressed with CYP11A in the zona fasciculata and zona reticularis, bovine CYP11B preferentially makes 11β-hydroxylated products. When bovine CYP11B is expressed alone in the zona glomerulosa of the adrenal cortex, it preferentially makes 18-hydroxylated products. However, this does not change the primary result that CYP11B2 must maintain both functions to preserve the conserved physiological configuration of the corticosteroid products.

Nonetheless, the functional results presented here should be viewed in the light that allosteric regulation by CYP11A would modify both 11- and 18-oxidation activities in the ancestral enzymes. There is some evidence that the human CYP11B paralogs are not allosterically regulated by CYP11A (Cao and Bernhardt 1999) suggesting the allosteric regulation by CYP11A may have been lost after the duplication of CYP11B in the primate lineage. It is therefore probable that their contemporaneous AncCYP11As regulated primate AncCYP11B and possibly AncCYP11B1. Specifically, I expect the AncCYP11B to have higher 11β-hydroxylase and lower 18-oxidation activity when co-expressed with its ancestral CYP11A. It may also be that AncCYP11B1 had higher 11β-hydroxylase activity if it had not lost allosteric regulation by CYP11A by the first speciation node after the duplication. However, the functional dissection of AncCYP11B2 is not subject to this problem because its function in the zona *zona* glomerulosa was never altered by CYP11A as they are not co-expressed. Thus, the dissection of AncCYP11B2 is not subject to this caveat.

When tracing the effects of substitutions on the CYP11B2 lineage, all four substitutions affect protein function, though there is some non-significant background-dependence in their effects. They combine to ultimately produce a CYP11B2 enzyme with higher 18-oxidation that still retains 11β-hydroxylation activity. Both human CYP11B paralogs are capable of oxidation on both the 11β- and 18-carbons of 11-deoxycorticosterone (DOC). The proportion of 11β-hydroxylation to 18-oxidation activity by one enzyme may partially depend on how long the substrate is retained in the active site to undergo subsequent oxidations (Imai, Yamazaki, and Kominami 1998), which suggests activity could be modulated by subtle shifts in the overall protein structure.

The imbalance of mineralocorticoids or glucocorticoids can lead to a variety of diseases in humans (Marques, Silverman, and Sternberg 2009; Peter, Dubuis, and Sippell 1999; Connell, Fraser, and Davies 2001) and during the evolution from AnceCYP11B to AncCYP11B1 or AncCYP11B2, it would be critical to maintain the balance of these hormones. The combinations NEqG and NdLG have higher 11-hydroxylation activity than any other variant, but it is difficult to speculate without knowing the mutational trajectory from AncCYP11B to AncCYP11B1 or without knowing its cellular expression or dependency on CYP11A. Those variants also have much lower 11-hydroxylase activity than the extant humanCYP11B1. When tracing the path of change from Anc11B to Anc11B2, nearly all mutational trajectories are open in the sense that there is not a complete loss of either function. The exception is NEqG which has much lower 18-oxidation activity than the AncCYP11B. In humans, a deficiency in aldosterone leads to hypoaldosteronism, hyponatremia, hyperkalaemia, fatal salt loss, and failure to thrive (Peter, Dubuis, and Sippell 1999). Although aldosterone likely plays a similar role in extant primates, again, it is unknowable if the NEqG variant would have been deleterious without additional pathway context. The reconstruction of the contemporaneous CYP11A with the ancestral CYP11B proteins is needed to further understand how allosteric regulation interacts with the resolution of the duplication of CYP11B in primates.

Our results suggest a complex evolutionary history of gene duplication and functional evolution with the interplay of enzyme activity within the enzymes studied. We still do not understand the relative importance and interplay of these factors during evolution of this gene family or how common these factors are generally during the evolution of biochemical pathways. However, this study suggests that biochemical networks can be shaped by mechanistic constraints of the enzyme and tissue-specific regulation with selection to avoid deleterious pleiotropy.

## Acknowledgement and funding sources

I want to thank J. W. Thornton and all members of the Thornton lab for their comments on this manuscript and support that enhanced this work. I especially want to thank L. Picton and A. Venkat for their support. Thanks to Michael Waterman for crucial plasmids. I want to thank A. Manning, K. Ferris, and C. Smukowski-Heil for their helpful comments on this manuscript. This work was supported by the National Institutes of Health (NRSA 1 F32 GM112350-01, C.F. O-M.) and initial funding from the National Institutes of Health (NIH) grant (R01-GM104397; J.W.T.)

