## Supplementary information for "Elaboration of the corticosteroid synthesis pathway in primates through a multi-step enzyme"

Supplemental information

Table S1

**Primers for human ferrodoxin reductase (FdxR), ferrodoxin (Fdx) and Cytochrome P450 11B (CYP11B) paralogs**

| **Primer target** | **RE** | **Sequence** |
| --- | --- | --- |
| Human FdxR Forward | NdeI | CGCCATATGAGCACCCAGGAAAAAACCCCGCAGATCTGTGTGG |
| Human FdxR Reverse | KpnI | GCGGGTACCTTAGCGCAGCATCTCCTGAG |
| Human Fdx Forward | NheI | CGCGCTAGCATGAGCAGCTCAGAAGATAAAATAACAGTCC |
| Human Fdx Reverse | HindIII | GCGAAGCTTAGGAGGTCTTGCCCACATCAATGGATTG |
| Human FdxR Forward | NdeI | CGCCATATGagcACACAGGAGAAGACCCCCCAGATCTGTGTGG |
| Human CYP11B1  Forward | NdeI | CGCCATATGGCTACTAAAGCTGCTCGTGTTCCACGTACAGTGCTGCCA |
| Human CYP11B1  Reverse | HindIII | GCGAAGCTTAATGATGATGATGATGATGGTTGATGGCTCTGAAGGTGAGGAG |
| Human CYP11B2  Forward | NdeI | CGCCATATGGCTACTAAAGCTGCTCGTGCCCCTAGGACGGTGCTGCCG |
| Human CYP11B2  Reverse | HindIII | GCGAAGCTTAATGATGATGATGATGATGGTTAATCGCTCTGAAAGTGAGGAG |

Table S2

Mass Spectronomy conditions

| **Compound** | **Parent ion (g/mol)** | **Primary ion fragment (g/mol)** | **Fragmentor energy (mV)** | **Collision energy (eV)** |
| --- | --- | --- | --- | --- |
| cortisol (standard) | 363.2 | 121.1 | 135 | 30 |
| aldosterone | 361.2 | 343.1 | 135 | 15 |
| corticosterone | 347.2 | 329.2 | 135 | 15 |
| DOC | 331.2 | 109.0 | 135 | 15 |

Supplemental Figures

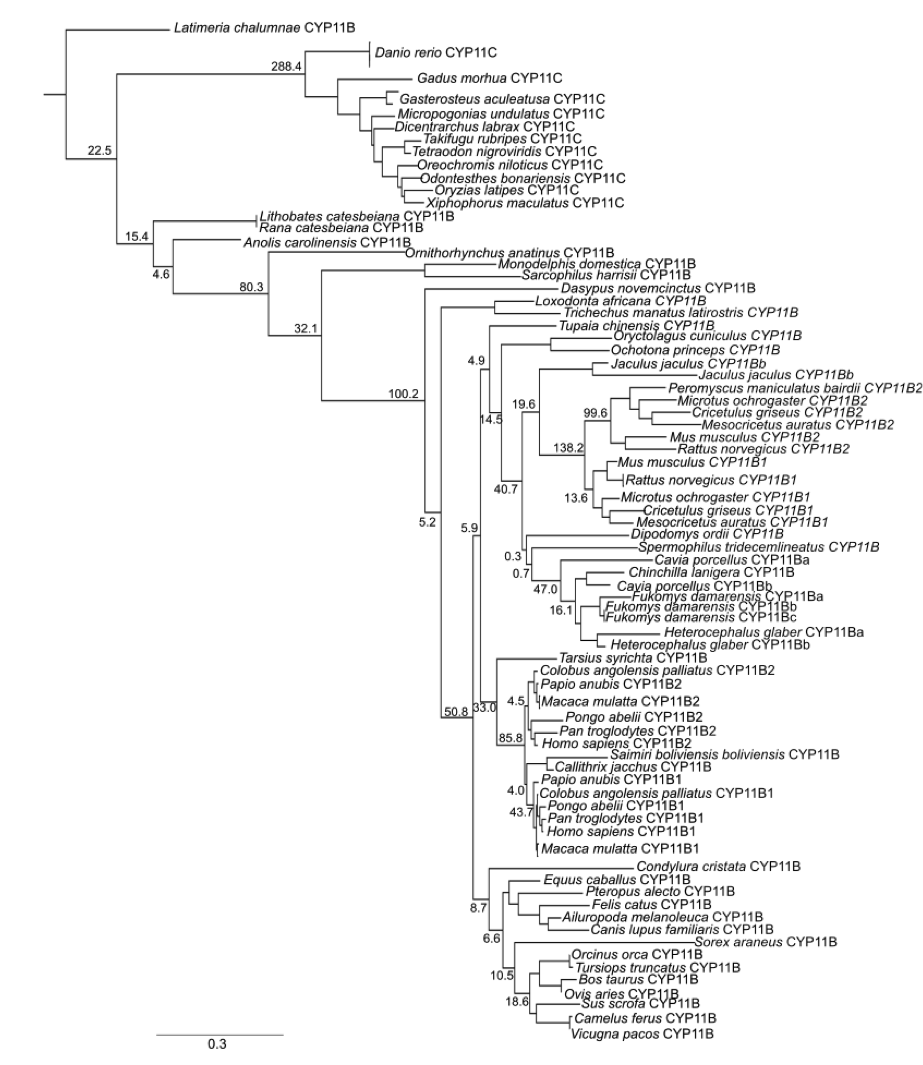

Figure S1 Maximum likelihood phylogeny of 86 vertebrate CYP11B/C sequences. The node support is calculated with the approximate likelihood ratio statistic.

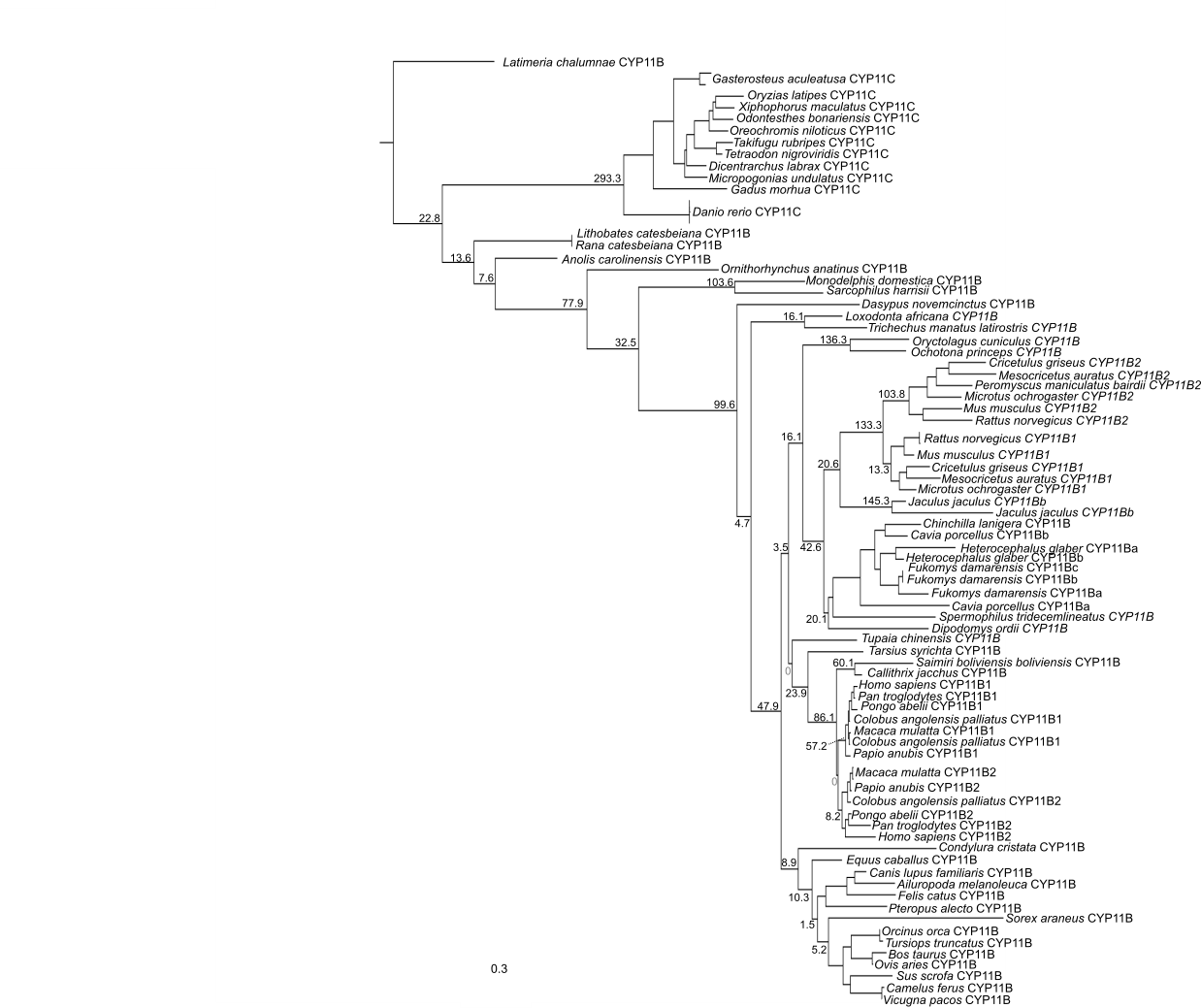

Figure S2 Maximum likelihood phylogeny with constrained nodes to conform to the known mammalian phylogeny labeled with light grey zeros as node support.

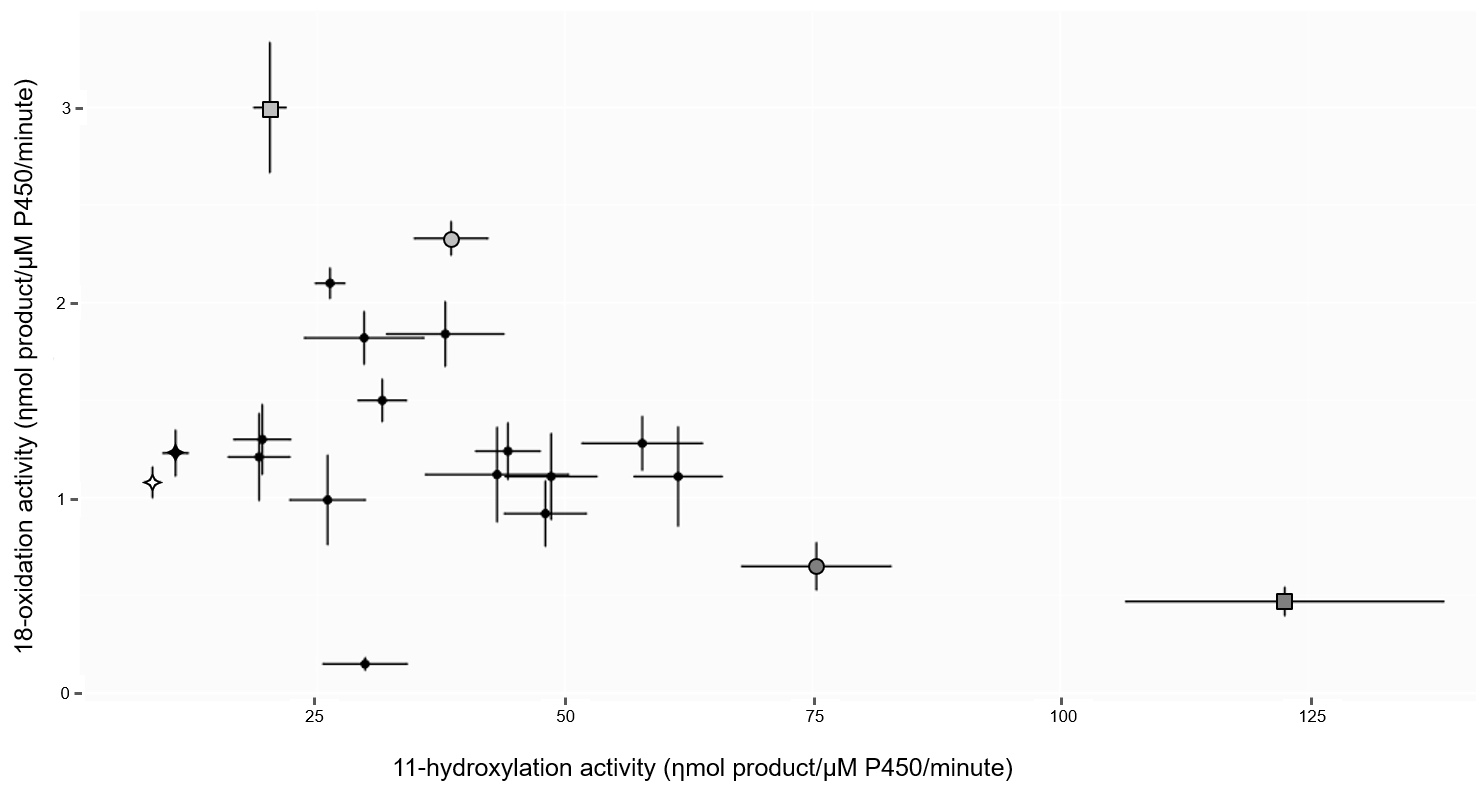

Figure S3 Activity of all CYP11B enzymes tested, including the intermediates (black circles) between AncCYP11B (black star) and AncCYP11B2 (dark grey circle). All other shapes, colors, and confidence intervals are the same as Figure 1 in the main text.
